# Simple cloning of large natural product biosynthetic gene clusters from Streptomyces by an engineered CRISPR/Cas12a system

**DOI:** 10.1101/2020.06.25.170191

**Authors:** Mindong Liang, Leshi Liu, Weishan Wang, Xiaoqian Zeng, Jiakun Liu, Loganathan Karthik, Guoliang Zhu, Linquan Bai, Chengjian Hou, Xiangyin Chen, Liming Ouyang, Xueting Liu, Bin Hu, Xuekui Xia, Yaojun Tong, Chunbo Lou, Gil Alterovitz, Gao-Yi Tan, Li-Xin Zhang

**Affiliations:** State Key Laboratory of Bioreactor Engineering (SKLBE), East China University of Science and Technology (ECUST), Shanghai 200237, China; State Key Laboratory of Microbial Resources, Institute of Microbiology, Chinese Academy of Sciences (CAS), Beijing 100101, China; Institute of Synthetic Biology, Shenzhen Institutes of Advanced Technology, Chinese Academy of Sciences (CAS), Shenzhen 518055, China; School of Life Sciences and Biotechnology, Shanghai Jiao Tong University (SJTU), Shanghai 200240, China; Institute of Animal Science, Guangdong Academy of Agricultural Sciences, Guangzhou 510640, China; Key Biosensor Laboratory of Shandong Province, Biology Institute, Qilu University of Technology (Shandong Academy of Sciences), Jinan, 250013, China; Boston Children’s Hospital, 300 Longwood Ave, Boston, MA 02115, USA

**Keywords:** CRISPR/Cas12a, natural product, simple cloning, biosynthetic gene cluster, large DNA fragment

## Abstract

Directly cloning of biosynthetic gene clusters (BGCs) from microbial genomes has been revolutionizing the natural product-based drug discovery. However, it is still very challenging to efficiently clone, for example, large (> 80kb) and GC-rich (> 70%), streptomycete originating BGCs. In this study, we developed a simple, fast yet efficient and low-cost *in vitro* platform for direct cloning large BGCs from streptomycete genomic DNA, named as CAT-FISHING (<u>C</u>RISPR/Cas12a- and <u>A</u>garose plug-based sys<u>T</u>em for <u>F</u>ast b<u>I</u>o<u>S</u>ynt<u>H</u>et<u>I</u>c ge<u>N</u>e cluster clonin<u>G</u>), by combining the advantages of CRISPR/Cas12a cleavage and bacterial artificial chromosome (BAC) library construction. CAT-FISHING was demonstrated by directly cloning large DNA fragments ranging from 47 to 139 kb with GC content of > 70% from the *S. albus* J1074 genome in a relatively efficient manner. Moreover, surugamides, encoded by a captured 87-kb BGC with GC content of 76%, was heterologously expressed in a *Streptomyces* chassis. These results indicate that CAT-FISHING is a powerful platform for BGCs batch cloning, which would be greatly beneficial to the natural products-based drug discovery. We believe that this system will lead a renaissance of interest in microorganisms as a source for drug development.

## 1. Introduction

Microorganisms, especially actinobacteria and fungi, remain unrivalled in their ability to produce bioactive small molecules (BSMs), some of which reached the market without any chemical modifications required, a testimony to the remarkable potential of *Streptomyces* to produce novel drugs. The expedition of *Streptomyces* genomes deciphered a large unexploited pool of novel biosynthetic gene clusters (BGCs), responsible for new but silent BSMs ^[1]^. However, the cloning of BGCs in *Streptomyces* is often very difficult because of the high GC and large size of those BGCs. It was found that 92% (1760/1910) of the characterized BGCs are smaller than 85kb, and 40% (756/1910) with > 70% GC content. In *Streptomyces*, 84% (534/634) BGCs have a GC content over 70% (Figure S1). To date, various processes have been developed for BGC cloning, such as genomic library (*i.e*., cosmid and fosmid) construction, recombination-based or RecET/Redαβ-based cloning, and Gibson assembly *etc* (Table 1). Additionally, the emergence of the CRISPR/Cas9 technique has enabled several new DNA cloning methods such as ExoCET, and CATCH *etc* ^[2–4]^. However, a simple, fast and efficient strategy for large BGCs, especially with high GC content, is still in urgent need to make natural BSMs accessible and affordable.

**Table 1.**
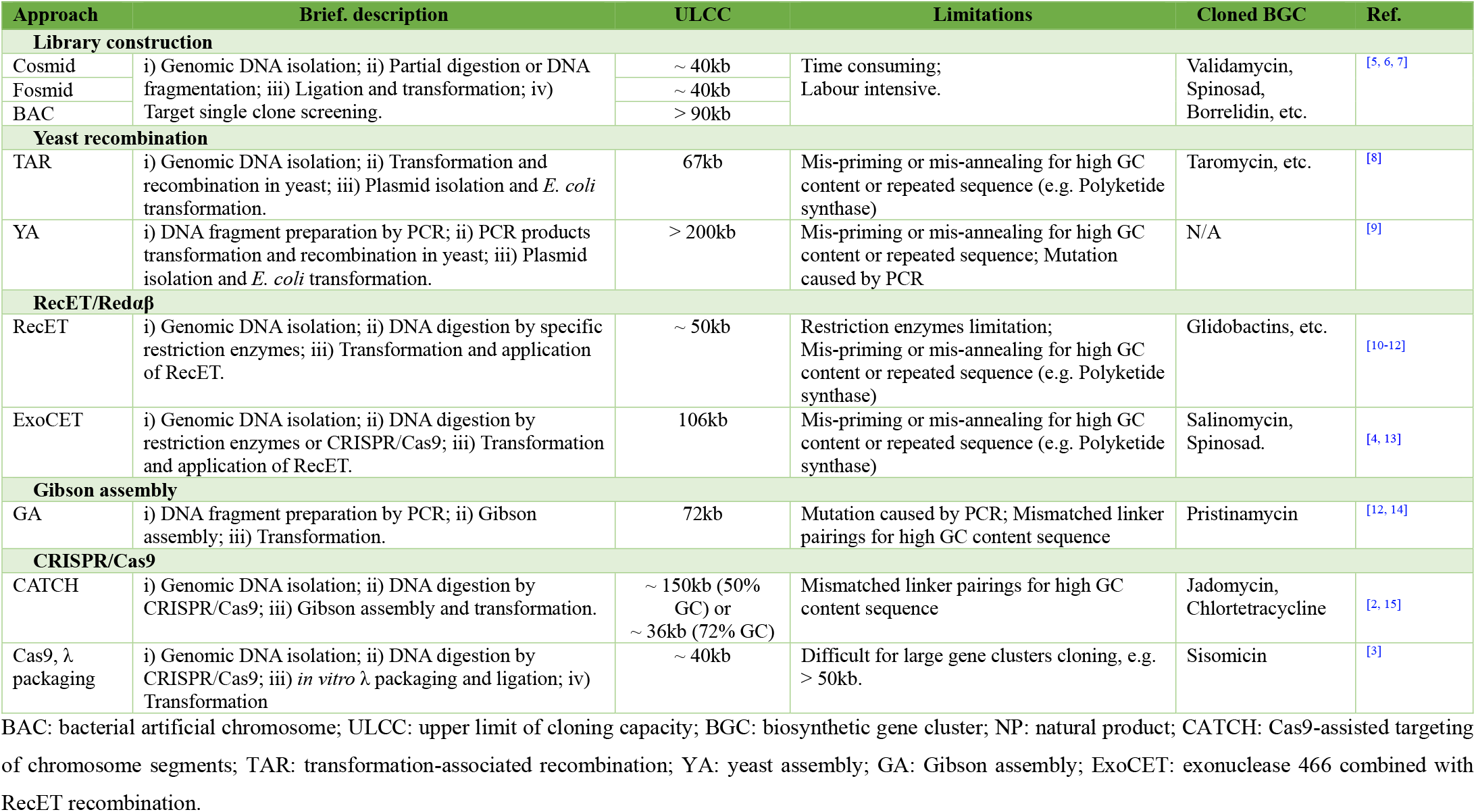
Previous approaches that have been used for BGC cloning *in vitro*.

CRISPR/Cas12a is a single RNA-guided (crRNA) endonuclease of a Class II CRISPR/Cas system ^[16]^. Unlike Cas9 proteins, Cas12a recognizes a T-rich protospacer-adjacent motif (PAM) instead of a G-rich PAM and generates dsDNA breaks with staggered ends instead of blunt ends. Besides the genome editing applications ^[16, 17]^, CRISPR/Cas12a has been widely used in nucleic acid-based diagnostic applications ^[18]^, small molecule detection ^[19, 20]^ *etc*. Moreover, it is worth noting that CRISPR/Cas12a possesses obvious superiority in DNA assembly with regard to its programable endonuclease activity and the DNA sticky ends of 4-or 5-nt overhangs ^[16]^. Based on these features, Li *et al*. developed a CRISPR/Cas12a-based DNA assembly standard, namely C-Brick ^[21]^. It was further reported that the accuracy of CRISPR/Cas12a cleavage could be significantly improved by shortening the length of crRNA, based on this, a DNA assembly method CCTL (Cpf1-assisted Cutting and Taq DNA ligase-assisted Ligation) was developed for efficient replacement of promoter region (< 1kb) in a 36-kb antibiotic BGC ^[22]^. And we foresee the potential of CRISPR/Cas12a reported by Lei et al might help to achieve the cloning of large DNA fragment with high GC content from genomic DNA sample. However, the susceptibility of large DNA fragments (> 80kb) to shearing in solution makes *in vitro* cloning of large BGC technically difficult, and how to achieve such a challenging objective by CRISPR/Cas12a was not clear.

As a classical method, by using agarose plug, bacterial artificial chromosome (BAC) library construction has been widely used for large DNA fragments cloning, but it is often time-consuming, labour intensive, expensive and technical demanding ^[5, 23]^. Though some new approaches of DNA cloning have been developed, for example, PCR-based cloning, Gibson assembly and recombination-based cloning, they often require additional homologous sequences among different DNA parts for assembly, which is relatively inefficient when meeting complicated DNA sequences (e.g. large fragment, high GC and/or highly repetitive) (Table 1). On the contrary, the BAC library is also indiscriminate toward insertion in a DNA sequence, this makes BAC library construction suitable for application to high GC content DNA samples.

In this study, by combining the DNA cleavage activity of CRISPR/Cas12a with the unique advantages of a BAC library construction, we have developed a CRISPR/Cas12a-and agarose plug-based method, designated CAT-FISHING (<u>C</u>RISPR/Cas12a- and <u>A</u>garose plug-based sys<u>T</u>em for <u>F</u>ast b<u>I</u>o<u>S</u>ynt<u>H</u>et<u>I</u>c ge<u>N</u>e cluster clonin<u>G</u>), which uses agarose plug-based *in situ* DNA isolation, digestion and liagtaion. As a proof of concept, large DNA fragments (or BGCs) of > 80kb have been fast cloned from *Streptomyces* genomic DNA (73% GC) by CAT-FISHING. Furthermore, the captured 87-kb surugamides BGC has also been successfully expressed in the *S. albus* J1074-derived cluster-free chassis strain ^[24]^.

## 2. Results

### 2.1. Design and workflow of CAT-FISHING for genome mining-based drug discovery

The flow chart of CAT-FISHING for capturing genome mining-based BGCs for drug discovery is presented in Figure 1. Mining microbial resources (*e.g*., *Streptomyces*) with advanced genome sequencing and bioinformatics provides new opportunities for natural product-based drug discovery. By using CAT-FISHING, target BGCs can be captured only with three steps. The first step is the capture plasmid construction and CRISPR/Cas12a-based plasmid digestion. In this step, two homolog arms (each arm containing at least one PAM site) that flank the target BGC were selected as adapter sequence and amplified by PCR. Then the BAC plasmid backbone containing the two adapter sequences and selection marker (*e.g*., antibiotic resistance gene, counter selection gene or *lacZ*), and the designated capture plasmid was constructed via the DNA assembly method. Under the guidance of crRNAs, the two selected PAM motif regions on the left and right adapter sequence were simultaneously digested by CRISPR/Cas12a, resulting in the linear capture plasmid. The second step is agarose plug-based genomic DNA isolation and *in situ* CRISPR/Cas12a digestion. According to the BAC library construction protocol^[7]^, genomic DNA plugs from the target strain were prepared. And the genomic DNA was digested *in situ* by the CRISPR/Cas12a system guided by the two designed crRNAs that were previously used in step one. The last step is ligation, desalting and transformation. The resulting linear capture plasmid and the digested genomic DNA from steps I and II, respectively, were gently mixed and ligated by T4 DNA ligase. Then desalted ligation products were introduced into *E. coli* by electroporation. The successfully captured BGCs can then be identified by a PCR screening approach. Lastly, the encoded compound of the captured target cryptic BGC can be obtained by heterologous expressing it in a suitable host with further chemical identification.

**Figure 1.**
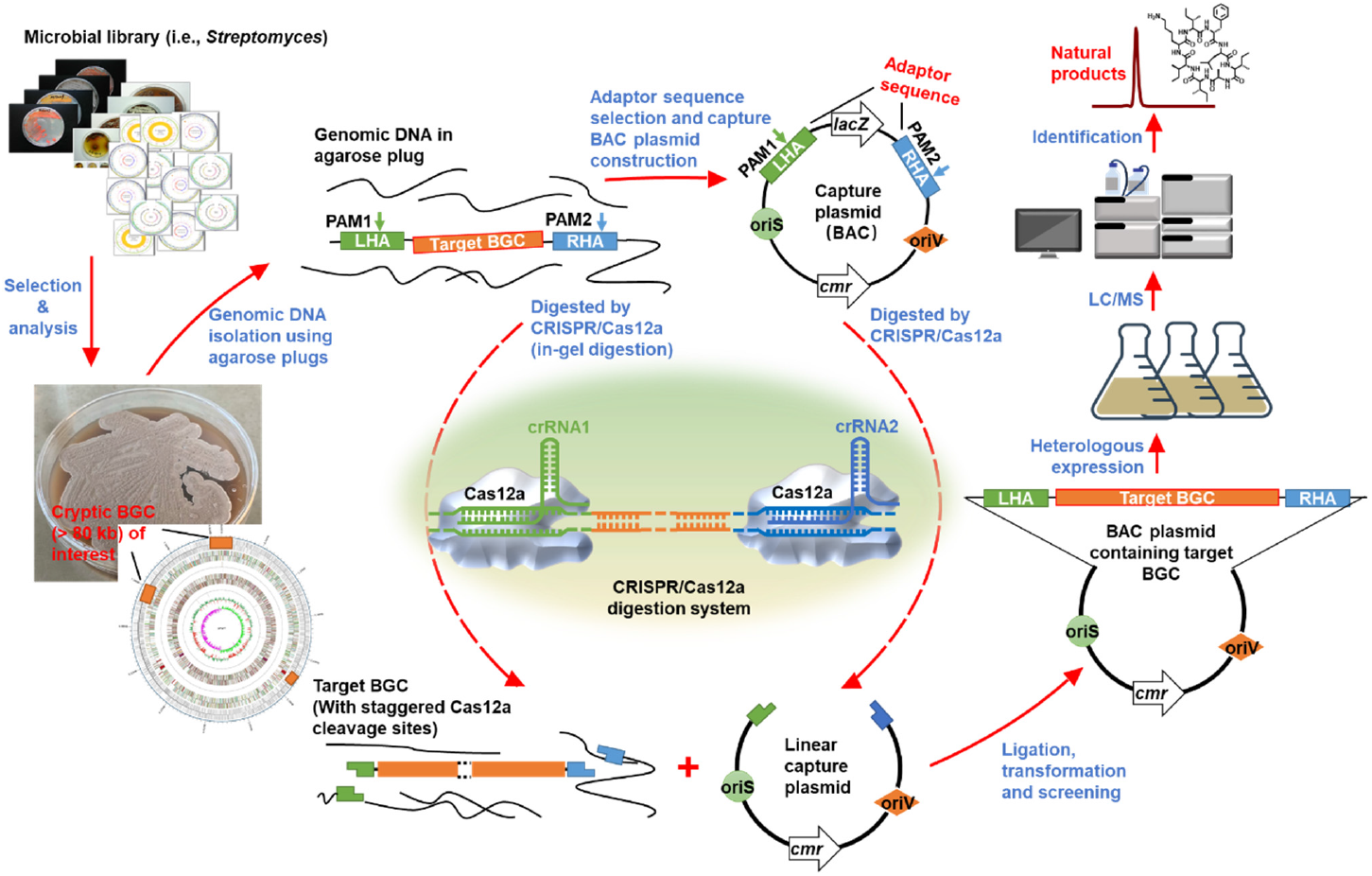
Schematic diagram of CAT-FISHING application. LHA: left homology arm, RHA: right homology arm, BGC: biosynthetic gene cluster; PAM: protospacer adjacent motif.

### 2.2. Evaluation of CRISPR/Cas12a-based DNA cloning efficiency

As CCTL, the principle underlying CAT-FISHING is the cohesive end ligation of two linearized DNA fragments by T4 DNA ligase. However, different from the widely used restriction endonuclease-based DNA cloning methods, here the cohesive ends were generated by paired crRNA-guided CRISPR/Cas12a cleavage. Though our previous study demonstrated that CRISPR/Cas12a cleaves more specifically on target DNA with a shorter spacer (*e.g*., 18nt) ^[22]^, the Cas12a cleavage is still not as accurate as NEB restriction endonucleases do. It is well known that the successful rate of direct cloning of the targeted large DNA fragment is often determined by the cloning efficiency. This study therefore carefully evaluated and compared the cloning efficiencies achieved by applying two different kinds of cohesive ends that were individually generated by NEB restriction endonuclease and CRSIPR/Cas12a.

As shown in Figure 2A, plasmid pGY2020 derived from the pCC2-FOS Fosmid vector (Epicentre) was constructed, and this plasmid contains two PAM sites (PAM1 and PAM2) as well as two NEB restriction endonucleases (EcoRI and HindIII). And the specific DNA fragment (ampicillin resistant gene Amp^R^, < 1kb) was cloned into pGY2020 by the CRISPR/Cas12a or NEB restriction enzymes-based method. There was no significant difference (P > 0.05) in the true positive rate between these two cloning methods (Figure 2B; Figure S2). In addtion, three randomly selected clones containing pGY2020/P1P2 also confirmed by junction sequencing and found no mutations (Figure 2C-D). These results indicate that CRISPR/Cas12a-based cloning strategy is able to clone short DNA fragments with a relatively high efficiency. Next, we would like to assess the ability of CRISPR/Cas12a-based method in direct cloning of large DNA fragments.

**Figure 2.**
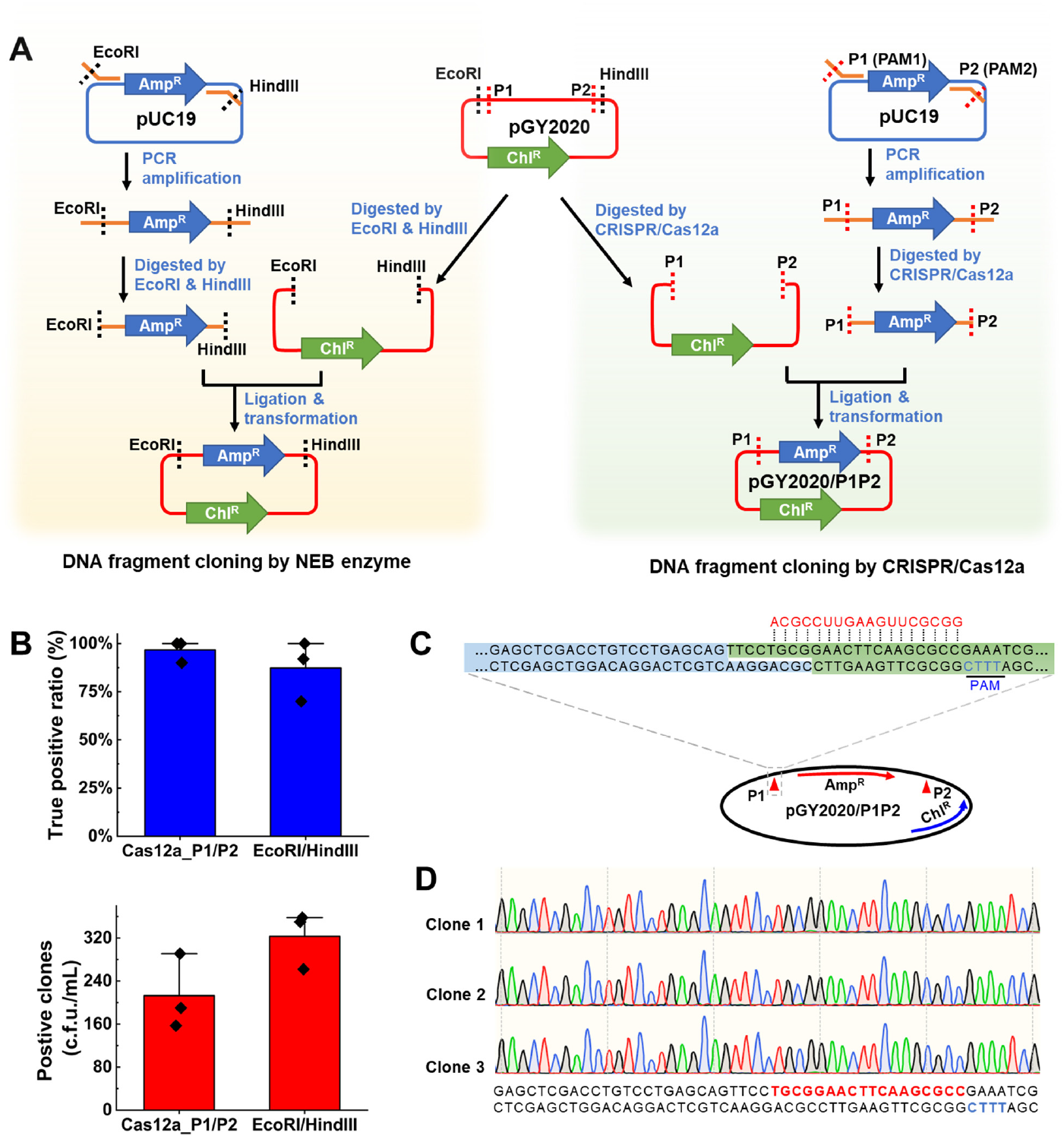
Evaluation and comparison of CRISPR/Cas12a-based and NEB restriction enzymes-based DNA cloning efficiency. A) Workflow of DNA fragment cloning by the CRISPR/Cas12a or NEB restriction enzymes-based method. B) Comparison of the clone numbers and positive rates of the CRISPR/Cas12a-based and NEB restriction enzymes-based methods. The results form three repeats are shown. C) Schematic representation of crRNA design and cohesive end ligation. 18-nt spacer crRNAs were employed, Cas12a mainly cleavage after the 14th base, generating 8-nt cohesive ends ^[22]^. D) Conformation of Cas12a-mediated cohesive end ligation by junction sequencing. Three clones were randomly selected for sequencing..

### 2.3. Cloning of a target DNA fragment from a BAC plasmid

In order to further demonstrate CRISPR/Cas12a-based cloning strategy in a simplified system, an 137-kb BAC plasmid pBAC-ZL (68% GC) was used to mimic and evaluate its cloning performance on a large DNA fragment. As shown in Figure 3A, a 50-kb fragment and an 80-kb fragment could be obtained by using the corresponding crRNAs-guided CRISPR/Cas12a cleavage. Under the guidance of the corresponding crRNA pairs, the BAC plasmid pBAC-ZL was digested by CRISPR/Cas12a, and 50-kb and 80-kb target bands were observed on the agarose gel after PFGE (pulsed field gel electrophoresis) (Figure 3B). By using the corresponding capture plasmid, two target DNA fragments were also successfully cloned from the BAC plasmid pBAC-ZL, as shown in Figure 3C–3E and Figure S3–S4. For 50-kb DNA fragment, of more than 100 transformants about 95% were the right clones. For the 80-kb DNA fragment, the number of transformants and the true positive rate were both lower, and about 50% of the transformants were the right clones. These results indicated that in a simplified system, by applying CRSIPR/Cas12a and corresponding capture plasmids, it could achieve high cloning rates of the 50-kb and 80-kb DNA fragments from the purified BAC plasmid DNA sample. It also needs to be noted that, for the 80-kb DNA fragment, the cloning difficulty was obviously greater.

**Figure 3.**
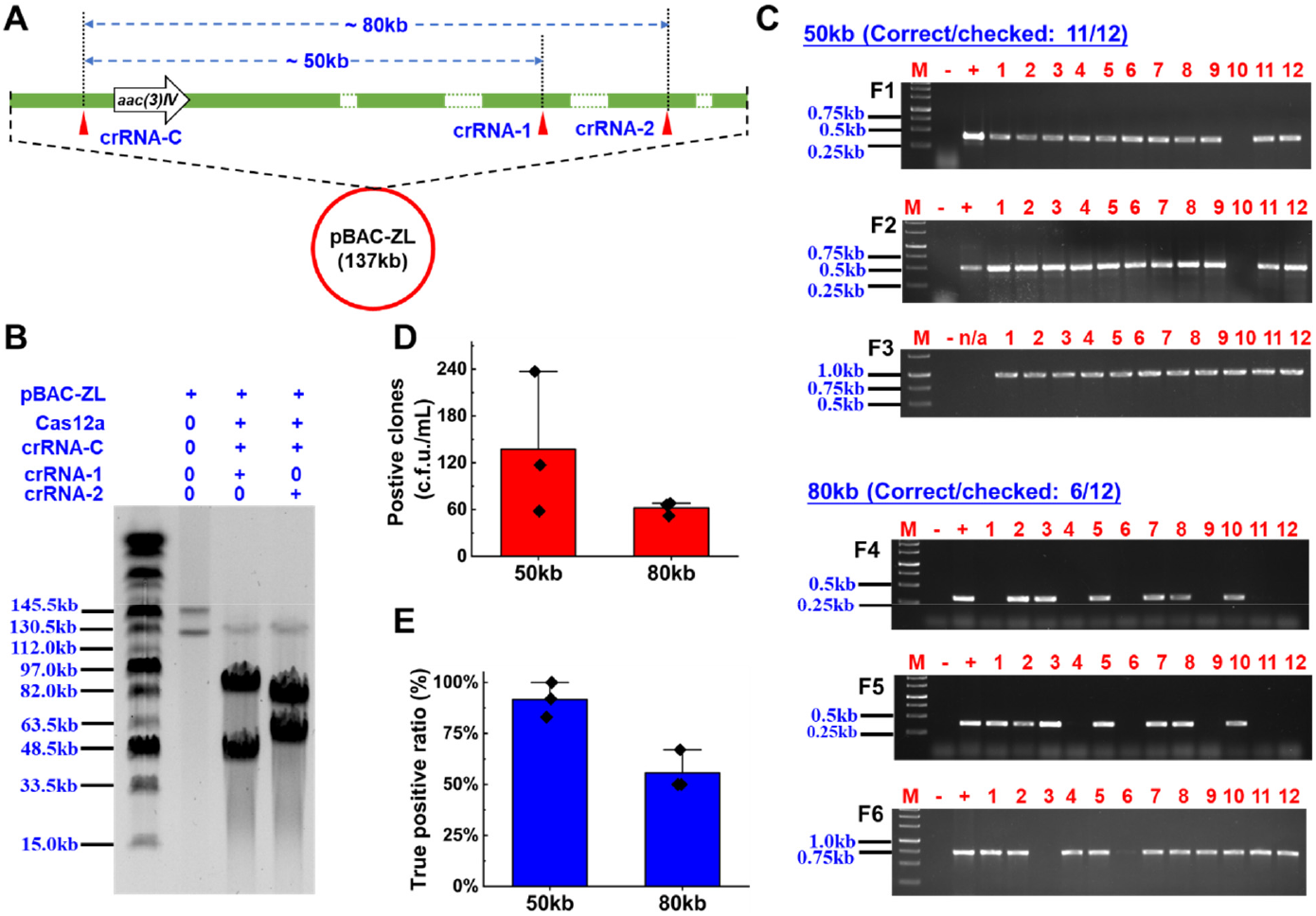
Direct cloning of 50kb and 80 kb of DNA fragments from pBAC-ZL by CRISPR/Cas12a-based cloning strategy. A) Two different target segments with different lengths (50 kb and 80 kb) in the BAC plasmid. B) Analysis of CRISPR/Cas12a-digested BAC plasmid (pBAC-ZL) by PFGE. PFGE was performed in 0.5% agarose at 6 V/cm with a 1 ~ 25 sec switching pulse time for 16 ~ 18 h in 0.5 × TBE buffer. C) PCR screening of right clones containing target DNA fragments (50kb or 80kb). Twelve clones were randamly selected from each agar plate. Three different primer pairs for regions in the middle and in the left/right boundary of target DNA fragments were used in the assay. F1, F2 and F3 are the PCR products that were amplified using 50-BAC-scr-up-F/R, 50-BAC-scr-middle-F/R and BAC-scr-down-F/R, respectively. F4, F5 and F6 are the PCR products that were amplified using 80-BAC-scr-up-F/R, 80-BAC-scr-middle-F/R and 80-scr-down-F/R, respectively. “-” represented blank control, genomic DNA of *E. coli* DH10B was used as PCR template. “+” represented positive control, pBAC-ZL plasmid DNA was used as PCR template. D-E) Determination of the clone numbers and positive rates for the DNA fragments of different lengths in three independent experiments/plates.

### 2.4. Direct cloning of target BGCs from *Streptomyces* genomic DNA by CAT-FISHING

In order to setup CAT-FISHING for directly cloning large BGCs from *Streptomyces* genomic DNA, CRISPR/Cas12a has been introduced into the initial process of BAC library construction by replacing the type II restriction enzymes (e.g. *Hind*III, *Eco*RI, *Bam*HI), which are involved in partial digestion ofthe genomic DNA (Figure 5). The optimized workflow of the target BGC capturing by CATFISHING is shown in Figure 4A. Based on our extensive investigation and test, we found that after genomic DNA isolation by agarose plugs and subsequent CRISPR/Cas12a *in situ* digestion, the resulting sample containing a mixture of genomic DNA could be directly used for subsequent ligation and transformation without prior DNA fragment isolation by PFGE and purification. This would allow CAT-FISHING to easily clone the large target DNA fragment.

**Figure 4.**
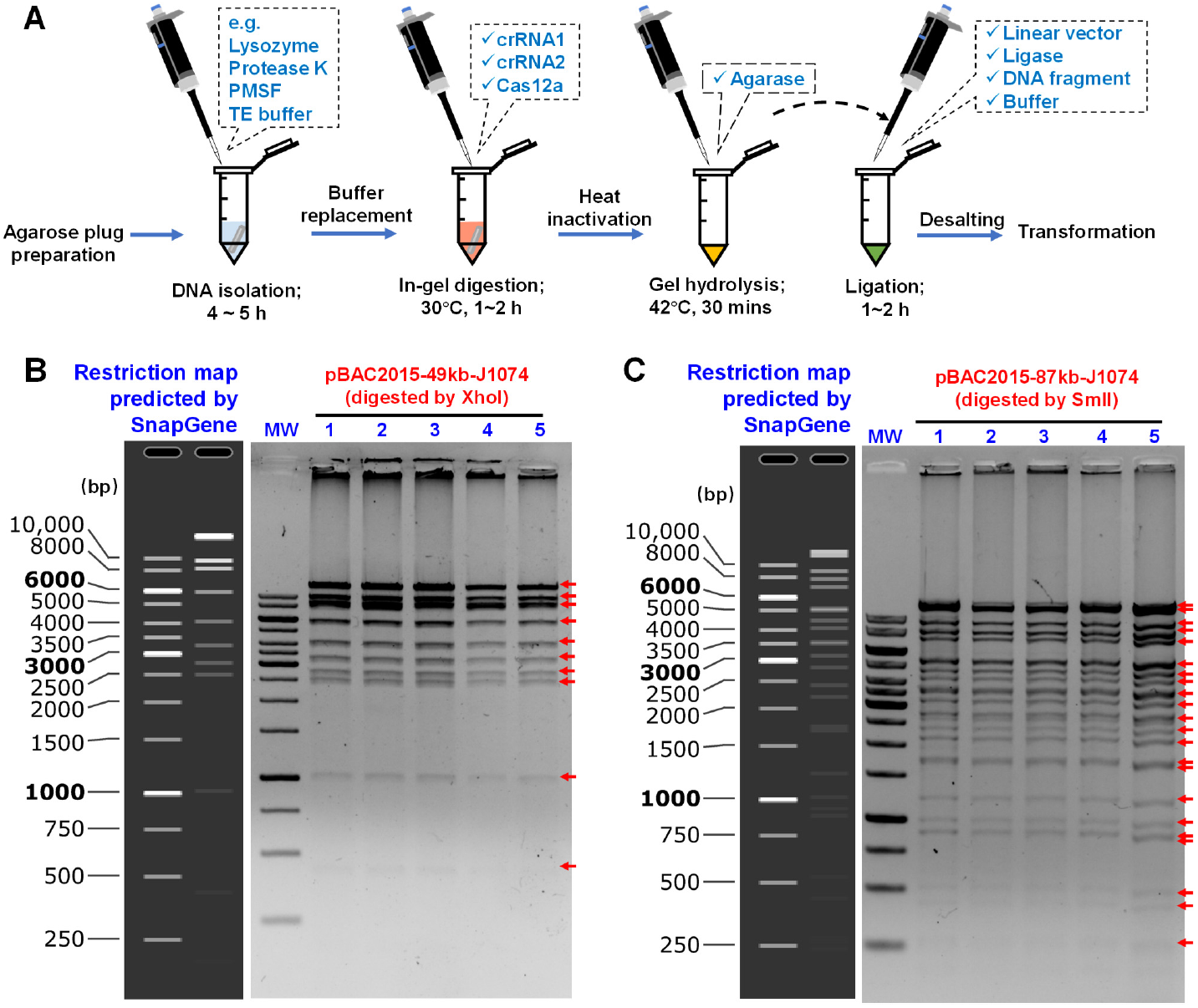
Direct cloning of large BGCs from genomic DNA by CAT-FISHING. A) Workflow of the target BGC capturing from genomic DNA by CAT-FISHING. B-C) Validation of five randomly selected positive clones containing a paulomycin or surugamides gene cluster by restriction enzymes digestion. XhoI and SmlI have, respectively, been used for paulomycin and surugamides gene cluster restriction. Bands are indicated by arrows. pBAC2015-49kb-J1074 digested by XhoI: 19776bp, 9843bp, 8402bp, 5856bp, 4108bp, 3270bp, 2749(2499)bp, 1011bp, 435bp; pBAC2015-87kb-J1074 digested by XhoI:13832(12760)bp, 9053bp, 7701bp, 6978bp, 5176(5040)bp, 4530bp, 4091bp, 3597(3497)bp, 3135bp, 2816bp, 2412bp, 2181bp, 1731(1681) bp, 1206bp, 1016bp, 930bp, 524bp,433bp,277(262)bp.

**Figure 5.**
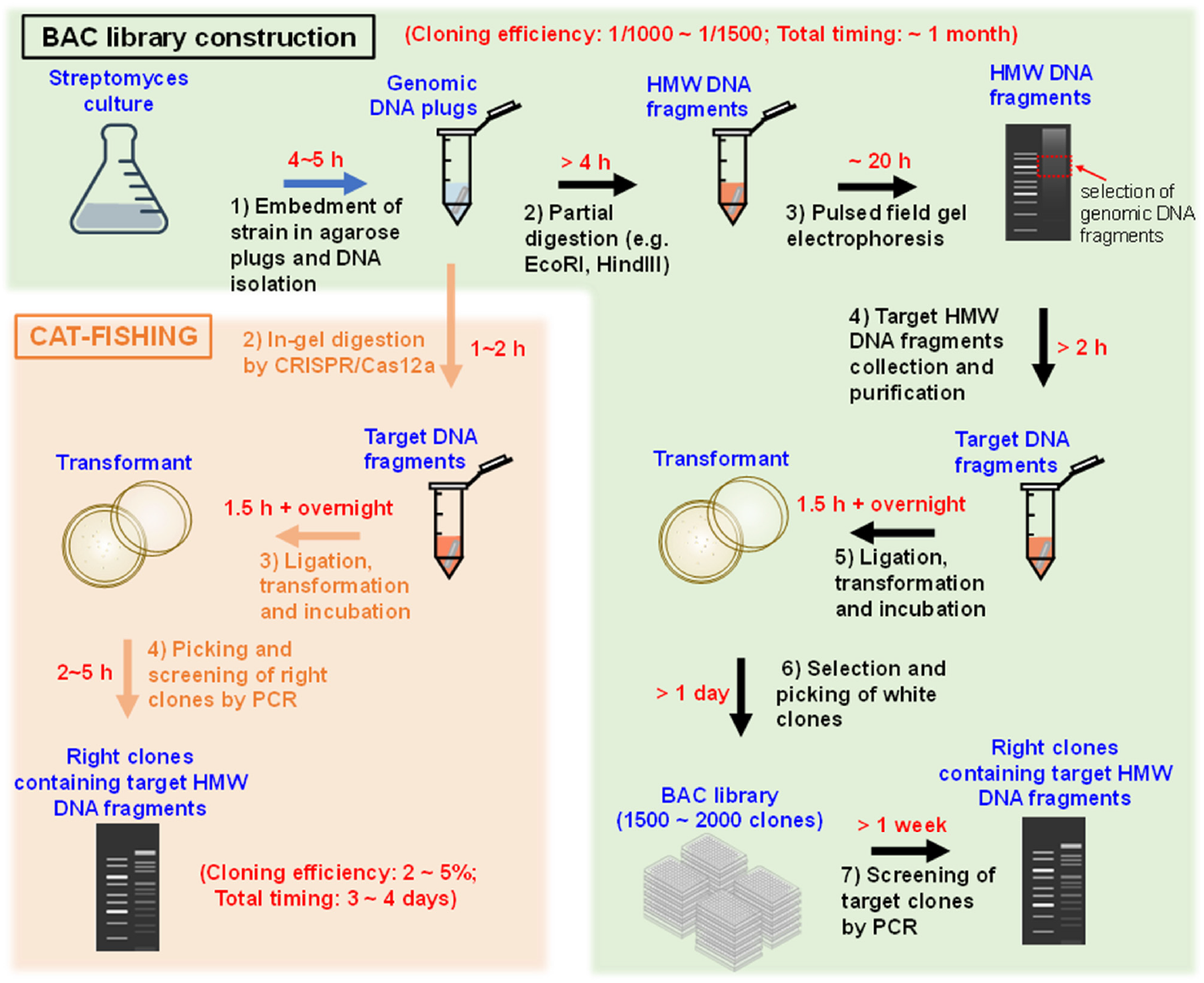
Methods of cloning large DNA fragments from genomic DNA by construction of BAC library (green box) or CAT-FISHING (orange box). HMW, High Molecular Weight.

In this study, a 49-kb paulomycin gene cluster ^[25]^, an 87-kb surugamides gene cluster ^[26]^ as well as a 139-kb candicidin gene cluster ^[27]^ were selected from the chromosome of *S. albus* J1074 to demonstrate our method. As shown in Table 2, the 49-kb paulomycin gene cluster (GC content 71%) and the 87-kb surugamides gene cluster (GC content 76%) were successfully cloned by CATFISHING, as confirmed by PCR and restriction mapping (Figure 4B–4C). And it was found the ratio of right clones that contain a 49-kb target BGC was 4~5%, and that for an 87-kb target BGC was 2~4%. Additionally, the 139-kb candicidin gene cluster (GC content 75%) was also cloned with CATFISHING, albeit with a much lower efficiency.

**Table 2.**
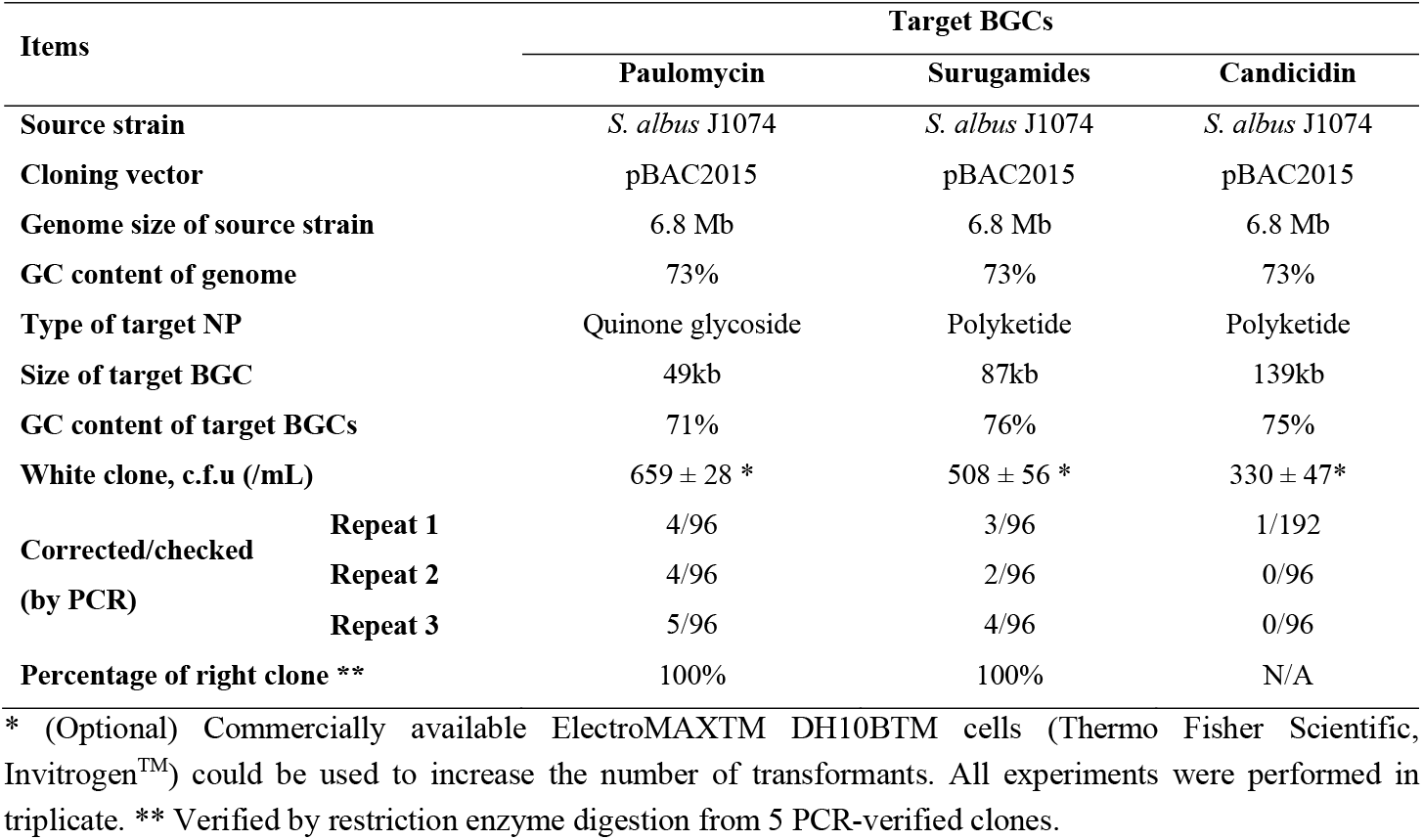
The cloning efficiencies of high-GC target BGCs from genomic DNA.

### 2.5. Expression of the target BGC in a cluster-free *Streptomyces* chassis strain

To thoroughly check the sequence and functional integrity of these BGCs that were captured by CATFISHING, as well as to prove access to genome mining through this route, a captured 87-kb surugamides gene cluster was expressed in a cluster-free *Streptomyces* chassis strain. The *aac(3)IV-oriT-attP(ΦC31)-int(ΦC31)* cassette was introduced into the target plasmid by Red/ET recombination. By applying ET12567/pUC307-mediated triparental conjugation, the resulting plasmid was integrated into the chromosome of *S. albus* Del14 (Figure 6A). During the subsequent fermentation study, surugamide A was produced in *S. albus* Del14-87kb by compared with *S. albus* Del14 and *S. albus* J1074 (Figure 6B). LC-MS/MS analysis further confirmed the production of surugamide A ([M+H]^+^ = 912.6252 Da, RT = 6.97 min) (Figure 6C).

**Figure 6.**
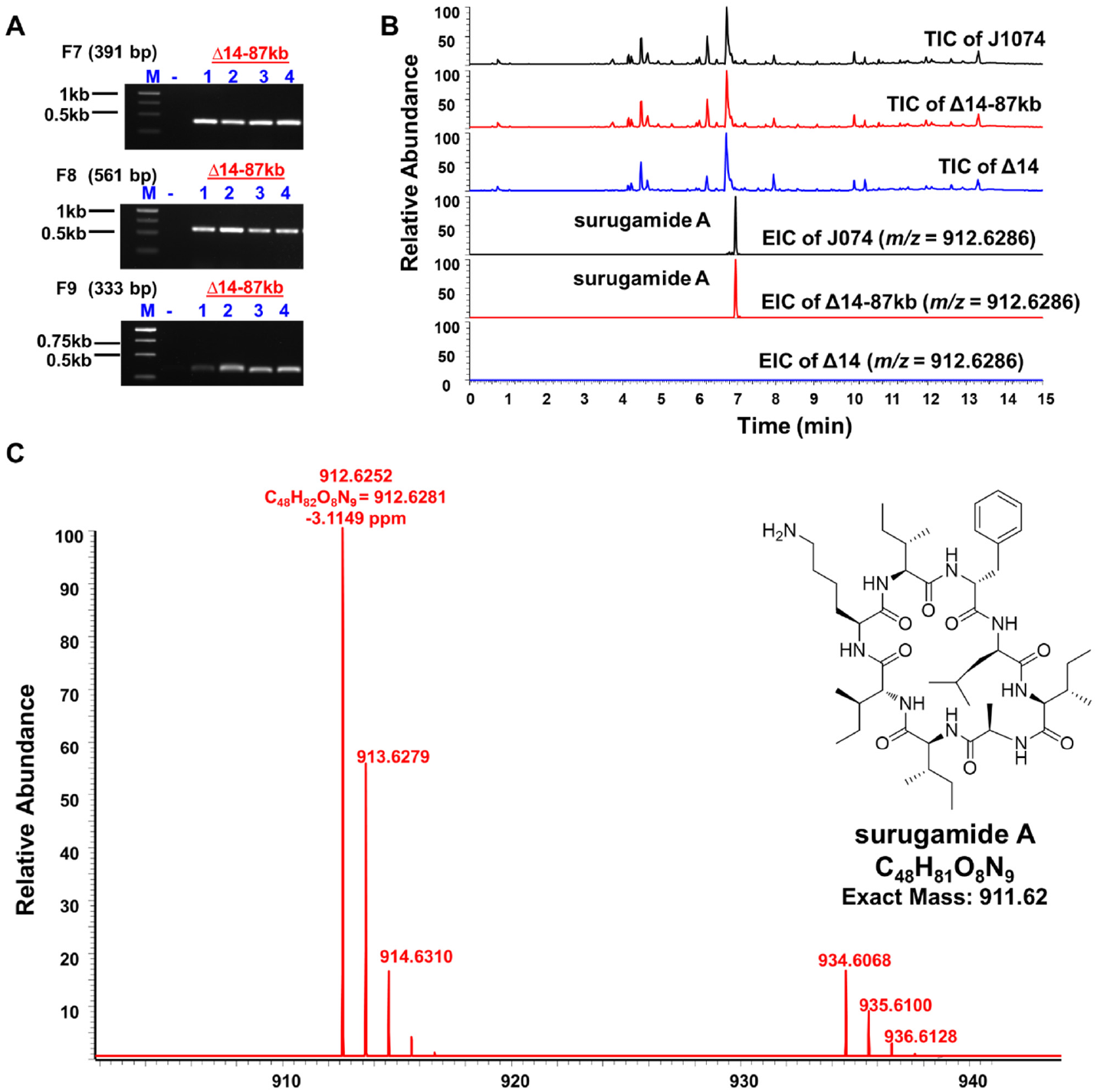
Expression of pBAC2015-87kb-J1074-int in *S. albus* Del14. A) PCR verfication of S. albus Del14-87kb. Four transformants were randomly slected. F7, F8 and F9 are the PCR products that were amplified using 87-scr-up-F/R, 87-scr-middle-F/R and 87-scr-down-F/R, respectively. “-” represented blank control, genomic DNA of *S. albus* Del14 was used as PCR template. B) Detection of surugamide components by LC-MS in S. albus Del-87kb. C) High-resolution mass spectrum of surugamide A ([M+H]+ = 912.6252 Da, RT = 6.97 min; [M+H]+ = 934.6068 Da, RT = 6.97 min).

### 2.6. The advantages of CAT-FISHING over other selected large DNA fragment cloning methods

The comparison between CAT-FISHING and BAC library construction as well as CAT-FISHING and other differents protocols of large DNA fragment cloning were presented in Figure 5, Table 3 and Table S1. From the comparison, it was concluded that CAT-FISHING is very simple, fast and efficient in cloning of large DNA fragment with high GC content. And the target large BGC could be obtained from *Streptomyces* samples in several days. In addtion, for high GC *Streptomyces* genomic DNA sample, the upper cloning limit of CAT-FISHING reached to as large as139 kb. Most importantly, by current cost accounting of each cloning steps, it was found that compared to BAC library construction, CAT-FISHING dramatically decreased the cost of *Streptomyces* originating target BGC cloning (Table 3; Table S2-S3). It makes that the CAT-FISHING is a totally affordable method for efficient and batch cloning large BGCs form *Streptomyces* or other microbial genomes.

**Table 3.**
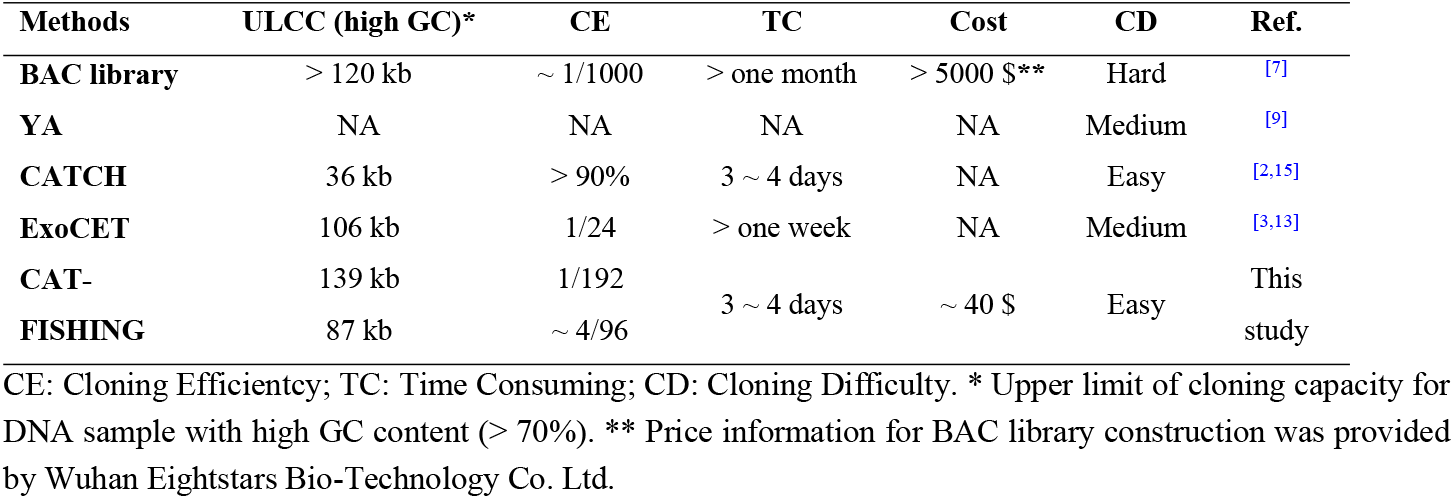
Comparison of selected methods of large DNA fragment (> 80kb) cloning.

| **Methods** | **ULCC (high GC)\*** | **CE** | **TC** | **Cost** | **CD** | **Ref.** |
| --- | --- | --- | --- | --- | --- | --- |
| **BAC library** | > 120 kb | ~ 1/1000 | > one month | > 5000 \$** | Hard | [7] |
| **YA** | NA | NA | NA | NA | Medium | [9] |
| **CATCH** | 36 kb | > 90% | 3 ~ 4 days | NA | Easy | [2,15] |
| **ExoCET** | 106 kb | 1/24 | > one week | NA | Medium | [3,13] |
| **CAT-** | 139 kb | 1/192 | 3 ~ 4 days | ~ 40 \$ | Easy | This study |
| **FISHING** | 87 kb | ~ 4/96 |  |  |  |  |

CE: Cloning Efficiency; TC: Time Consuming; CD: Cloning Difficulty. \* Upper limit of cloning capacity for DNA sample with high GC content (> 70%). \*\* Price information for BAC library construction was provided by Wuhan Eightstars Bio-Technology Co. Ltd.

## 3. Discussion

Cloning and manipulation of large DNA fragments is a fundamental and important platform technology for mining results BSMs from rich microbial genome sources. In order to avoid shearing of large DNA fragment during *in vitro* manipulation, in this study, CAT-FISHING combined the advantages of CRISPR/Cas12a cleavage and BAC library construction (agarose plug-based *in situ* DNA isolation and digestion). Instead of using restriction enzymes (*e.g*. HindIII, EcoRI, or BamHI) for the random fragmentation of the genomic DNA ^[7]^, CAT-FISHING utilizes CRISPR/Cas12a together with specific crRNA pairs to precisely *in situ* digestion of the genomic DNA in agarose gel (Figure 1; Figure 4A). This theoretically makes CAT-FISHING very suitable for cloning of large BGCs with high GC contents.

There are two concerns about CRISPR/Cas12a-based DNA cleavage, off-target (non-specific cleavage) and inaccuracy of the cleavage sites ^[21, 28]^. Both could reduce the cloning efficiency of CATFISHING. Lei *et al*. reported that the cleavage specificity of CRISPR/Cas12a would enhance with a shorter crRNA spacer (*i.e*., 17 ~ 19 nt) ^[22]^. In this study, we used crRNA with an 18-nt spacer to evaluate the cloning efficiency of CRISPR/Cas12a-based cloning strategy. As shown in Figure 2, the number of transformants obtained by CRISPR/Cas12a were fewer than the control, while no significant difference (*P* > 0.05) was observed. On the other hand, non-specific cleavage by CRISPR/Cas12a could be minimized by decreasing the Cas12a concentration, shortening the cleavage time and slection of more suitable crRNA. As a result, when a purified 137-kb BAC plasmid was used to test CRISPR/Cas12a-based cloning strategy, the plasmid was almost completely digested and no non-specific cleavage products appeared on the agarose gel (Figure 3A–3B). To some extent, these results verify the relatively higher efficiency of cloning 50-kb and 80-kb DNA fragments from BAC plasmid (Figure 3C–3E).

PFGE is a powerful and essential tool for isolating large DNA fragments. During BAC library construction, however, PFGE is time consuming, as it often takes 16 ~ 24 hours to separate specific size DNA fragments ^[7]^. Moreover, the following operational steps, such as DNA elution and purification (Figure 5), could drastically decrease the DNA integrity/amount as well as the subsequent ligation or transformation efficiency. In this study, after many preliminary tests, we found that, following agarose digestion, the resulting mixture from a CRISPR/Cas12a-treated genomic DNA plug could be directly used for ligation with the vector and subsequent electro-transformation (Figure 4A; Figure 5). Without the need for PFGE or the preparation of purified high molecular weight genomic DNA fragments, and compared to previously reported large DNA fragment cloning approaches (*e.g*. ExoCET, CATCH, TAR *etc*.), the cloning process in CAT-FISHING has been greatly simplified ^[2, 4, 8, 15]^.

In this study, by applying CAT-FISHING, we achieved a cloning efficiency of 2~4% when cloning an 87-kb target BGC (Figure 4B–4C; Table 2). It is close to Wang’s result that 106 kb salinomycin BGC from *S. albus* obtained by ExoCET at screen-positive rate of 4% ^[4]^. To further determine the upper cloning limit of CAT-FISHING, an 139-kb candicidin BGC was selected for demonstration. Although this study successfully cloned candicidin BGC from *S. albus* genome, the screen-positive rate of target clone was less than 1%. Probably due to the complexity of the un-purified DNA mixture sample, slightly inaccurate of Cas12a cleavage site^[22]^ and high activity of T4 DNA ligase, many short DNA fragments or incomplete pieces of target BGCs were inserted into capture plasmids. Therefore, it is reasonable to predict that, if necessary, through DNA fragment isolation and purification, the cloning performance toward a 139-kb BGC should also be dramatically improved. Anyway, compared to other methods of large DNA fragment cloning, including the BAC library, in which often only a few right strains could be screened out of thousands of clones (i.e., 1/1000 ~ 1/2000) ^[5, 29]^, CAT-FISHING is a simpler method with a greater efficiency for cloning target BGCs with high GC content.

*Streptomyces* are the source of a majority of antibiotic classes in current clinical and agricultural use^[30]^. *S. albus* J 1074 is one of the most widely used *Streptomyces* chassis for genome mining ^[31]^. In this study, *S. albus* Del14, which is a *S. albus* J 1074-derived cluster-free chassis strain ^[24]^, has been used to demonstrate the sequence and functional integrity of the BGCs obtained by CAT-FISHING. As shown in Figure 5, the 87-kb target BGC was successfully expressed, and the corresponding NP surugamides (inhibitors of cathepsin B) could be confirmed by LC-MS. During genome mining, BGC cloning and expression is the most important starting point for the next step of bioactivity analysis and structure identification of target BSMs. The current results present a case study for NP production in *Streptomyces* by applying CAT-FISHING. Lastly, as an *in vitro* manipulation platform, not limited to actinomyces, CAT-FISHING could easily be extended to fungi and other microbial resources ^[32]^.

As a results, we successfully combining the accurate DNA cleavage activity of CRISPR/Cas12a with the unique advantages of BAC library construction platform that is believed one of the most efficient strategies of cloning large DNA fragment with high G+C content. This combining gave the birth of CAT-FISHING, which possesses some unique features comparing to previous methods: 1) CATFISHING is a very simple and low-cost cloning method that can directly capture the target large DNA fragment (>80kb) with high GC content (70%) from genomic DNA; 2) CAT-FISHING is very fast, the target large BGC could be obtained from streptomycete genomic DNA samples in a few days (Table 3); 3) CAT-FISHING does not require PFGE or the preparation of purified large DNA fragments. Moreover, in addition to genome editing, DNA assembly, nucleic acid and small molecule detection *etc*., this study also expanded the application of CRISPR/Cas12a to direct cloning of large DNA fragments with complicated DNA sequence (e.g. high GC or sequences with repeat) *in vitro*. This innovation of a fundamental platform technology for use in genome mining through application of the CRISPR/Cas12a system would facilitate the discovery of novel BSMs from microbial sources.

## 4. Experimental Section

### Strains, plasmids and media

The strains and plasmids used in this work are present in Table S4. *Escherichia coli* and its derivatives were cultivated on Luria-Bertani (LB) agar plates (tryptone 10 g/L, yeast extract 5 g/L and NaCl 10 g/L, pH = 7.2). Streptomyces and its derivatives were cultivated on soybean flour-mannitol (SFM) agar plates (soybean flour 20 g/L, mannitol 20 g/L and agar 20 g/L, pH = 7.2) or ISP4 (International Streptomyces Project Medium 4, BD Bioscience, San Jose, CA, USA). In the fermentation experiments, seeds were grown in TSB (trypticase soy broth, Oxoid Ltd), and R4 medium was used for subsequent fermentations (in a 250-mL Erlenmeyer flask, 30°C and 200 rpm).

### Capture plasmid construction

The primers for capture plasmid construction are listed in Table S5. The capture plasmid was constructed by introducing the *lacZ* gene as well as two PCR-amplified homology arms (each arm containing at least one PAM site) corresponding to the flanking regions of the target DNA fragment (or BGC) in to pBAC2015 ^[10]^. Assembly of multiple DNA fragments was carried out by Gibson assembly or with the EZmax one-step seamless cloning kit (Tolo Biotechnology, Shanghai). Plasmid DNA was isolated from *E. coli* by using alkaline lysis ^[33]^.

### Genomic DNA isolation

For genomic DNA isolation, *S. albus* J1074 was cultured in Oxoid TSB (30 g/L) supplemented with glycine (5 g/L). According to *Practical Streptomyces Genetics* ^[33]^, after cultivation at 200 rpm and 30°C for 24 ~ 36 h, mycelium was collected by centrifugation (4°C, 4000 g, 5 min). Mycelium was resuspended in TE25S (25 mM Tris-HCl pH 8, 25 mM EDTA pH 8, 0.3 M sucrose) and then the supernatant was removed (4°C, 4000 g, 5 min). The mycelium density was adjusted with TE25S and it was mixed with an equal volume of 1.0% LMP agarose (1.0% molten solution of low melting point agarose) at 50°C, and then poured into holes in a plug mould (100-μl holes). The blocks were removed from the mould and incubated at 37°C for 1 h in lysozyme solution (2 mg/mL in TE25S). The lysozyme solution was removed and the blocks were incubated at 50°C for 2 h in proteinase K solution (1 mg/mL proteinase K in NDS. NDS: 0.5 M EDTA pH 8, 10 mM Tris-HCl pH 8, 1% SDS). The proteinase K solution was removed and the blocks were washed once for 30 min in TE (10 mM Tris-HCl pH 8, 1 mM EDTA pH 8) supplemented with 0.1 mM PMSF (phenylmethanesulfonyl fluoride, serine proteinase inhibitor), then three times with TE for 30 min. After removing all of the liquid, the agarose plugs could be used for CRISPR/Cas12a digestion, but could also be stored for up to 1 month at 4°C in 70% ethanol.

### crRNApreparation

The oligonucleotides used as templates for crRNA transcription are given in Table S6. According to our previous study ^[19]^, crRNA was prepared via *in vitro* transcription. Templates for crRNA synthesis were synthesized and annealed by using Taq DNA Polymerase PCR Buffer (Thermo Fisher Scientific). A HiScribe™ T7 Quick High Yield RNA Synthesis Kit (NEB) was used for crRNA *in vitro* transcription. The resulting crRNA was purified using RNA Clean & ConcentratorTM-5 kit (Zymo Research), and subsequently quantified using a NanoDrop 2000 spectrophotometer (Thermo Fisher Scientific). RNase-free materials (Axygen Scientific, Union City, CA, USA) and conditions were applied during the entire experimental process.

### CRISPR/Cas12a-based DNA restriction and ligation

The Cas12a (LbCas12a) protein used in this study was overexpressed in pET28a and then purified by fast protein liquid chromatography (FPLC; AKTA Explorer 100, GE Healthcare)^[19]^. In the CRISPR/Cas12a cutting system, NEBuffer™ 3.1 (100 mM NaCl, 50 mM Tris-HCl, 10 mM MgCl_2_, 100 μg/mL BSA, pH 7.9) was adopted as the reaction buffer. For pBAC-ZL or capture plasmid cleavage, plasmid DNA was incubated with Cas12a protein and the corresponding crRNA pairs at 37°C for 1 h. After the reaction, the linearized capture plasmids or DNA fragments of pBAC-ZL were prepared using isopropanol and ethanol ^[10]^. The resulting linear capture plasmid or DNA fragments of pBAC-ZL could be used for the following ligation. If necessary, the large DNA fragments could be analysed by pulsed field gel electrophoresis (PFGE) with the CHEFDR III apparatus (Bio-Rad, Richmond, CA). PFGE was performed in 0.5% agarose at 6 V/cm with a 1 ~ 25 sec switching pulse time for 16 ~ 18 h in 0.5 × TBE buffer. For genomic DNA cleavage, plugs were initially equilibrated in 1 × NEBuffer™ 3.1, then transferred into a cleavage system that contained the Cas12a protein and the corresponding crRNA pair, and finally incubated at 37°C for 1 ~ 2 h. After the reaction, following heat treatment at 65°C for 10 min to inactive Cas12a protein, the LMP agarose gel was hydrolysed using β-Agarase I (NEB) for 30 min at 42°C. Afterward, the resulting DNA mixture could be directly used for following the ligation with the corresponding linear capture vectors by T4 DNA ligase (NEB).

### Electro-transformation of E. coli

Following the transfer of ligation samples into 0.1 M glucose/1% agarose gel to desalt for 1 ~ 2 h on ice, these samples could be used for electro-transformation. The high efficiency electro-transformation of *E. coli* cells was accomplished according to a previous study^[34]^. The following electro-transformations were performed in 2-mm cuvettes using the Bio-Rad GenePulser Xcell™ system (electroporator conditions: 2500 V, 200 Ω and 25 μF). Then, to the *E. coli* cells in the cuvette, 1 mL of SOC medium (tryptone 20 g/L, yeast extract 5 g/L, NaCl 0.5 g/L, KCl 2.5 mM, MgCl_2_ 10 mM, glucose 20 mM) was added and the mixture was transferred into a 15-mL Falcom™ tube. After shaking at 200 rpm for 1 h at 37°C, the strains were collected and spread on selective LB agar plates. The plates were incubated overnight at 37°C, and the transformants were screened and verified by PCR using the primers listed in Table S7.

### Expression of BGCs in Streptomyces and LC-MS analysis

The *aac(3)IV-oriT-attP(ΦC31)-int(ΦC31)* cassette amplified from pSET152 has been introduced into BAC plasmids by Red/ET recombination^[35]^. The resulting plasmid was introduced into *S. albus* Del14 by triparental conjugation according to a previous study ^[5]^. The transconjugants were screened and verified by PCR using the primers listed in Table S7. After fermentation, the production of target natural product was qualitatively analysed using a high-resolution Q-Exactive Hybrid Quadrupole-Orbitrap mass spectrometer (Thermo Scientific, Waltham, MA).

## Supporting Information

Supporting Information is available from the Wiley Online Library or from the author.

## Acknowledgements

Mindong Liang, Leshi Liu, Weishan Wang and Xiaoqian Zeng contributed equally to this work. The authors would like to thank Prof. Dr. Fei Xu (School of Medicine, Zhejiang University) and Prof. Dr. Wei Ma (School of Life Sciences and Biotechnology, Shanghai Jiao Tong University) for their helpful suggestion in manuscript preparation and pulsed-field gel electrophoresis. This work was supported by the National Natural Science Foundation of China (31720103901, 31870040, 21877038), National Key Research and Development Project (2020YFA0907804, 2020YFA0907304), the “111” Project of China (B18022), the Fundamental Research Funds for the Central Universities (22221818014), the Shanghai Science and Technology Commission (18JC1411900), the Open Project Funding of the State Key Laboratory of Bioreactor Engineering, and the Shandong Taishan Scholar Award to L.-X. Z.

## ToC figure

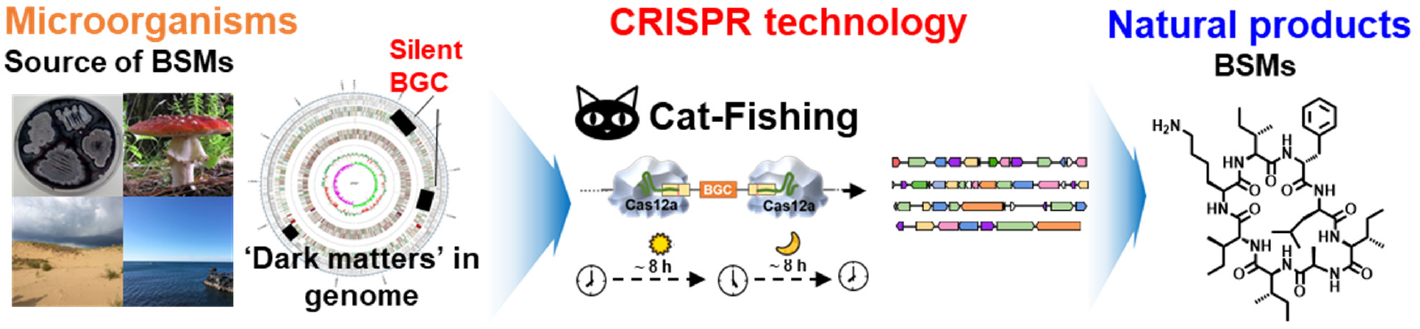
Directly cloning of large biosynthetic gene cluster (BGC) from even unculturable microbial genomes revolutionized nature products-based drug discovery. 47-139kb of target DNA fragments were simply and rapidly direct cloned from high GC genomic DNA sample by CAT-FISHING. Surugamides, encoded by a captured 87-kb gene cluster, was expressed and identified in a cluster-free Streptomyces chassis, validating the great potential of CAT-FISHING.

## Supporting Information

**Figure S1.**
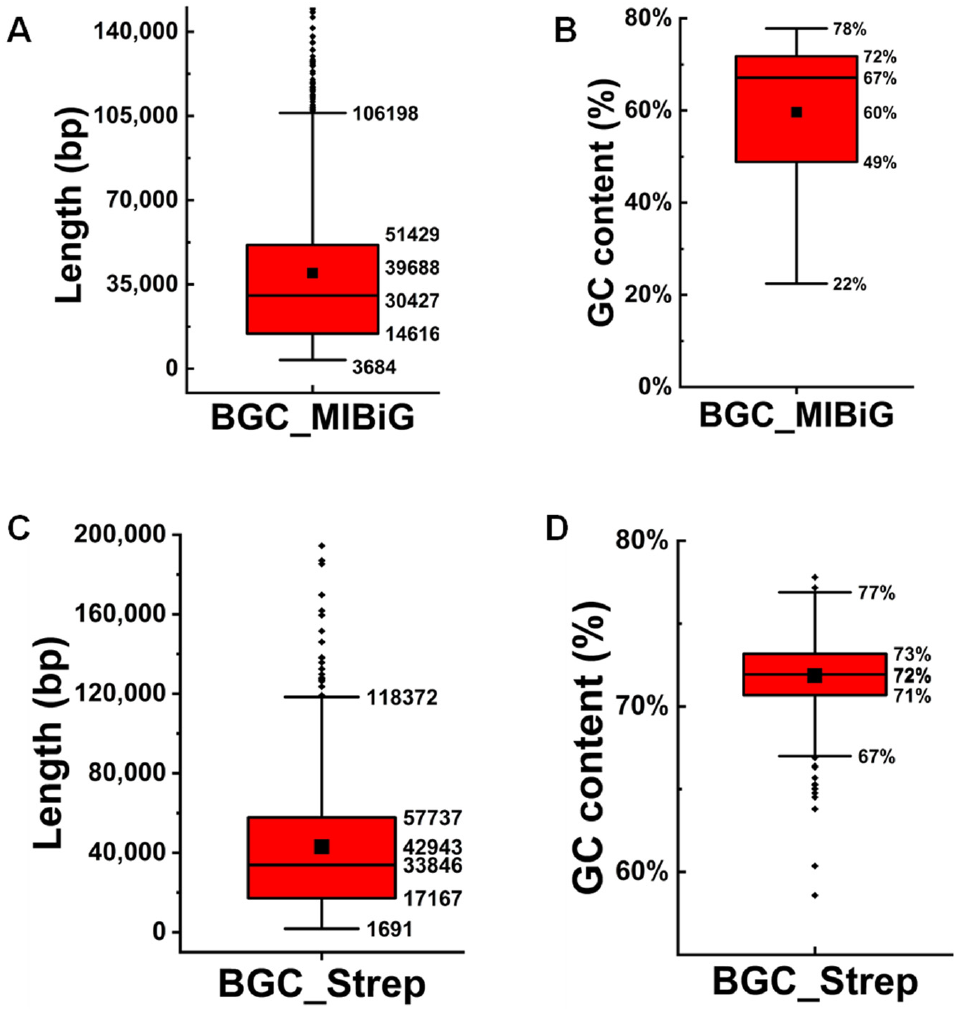
Statistical analysis of GC content and length of characterized BGCs in the MIBiG database. A-B. Distribution of BGCs length and GC content in characterized gene clusters identified in the MIBiG database. C-D. Distribution of BGCs length and GC content in characterized gene clusters identified in Streptomyces. 1910 characterized BGCs were download from MIBiG database (https://mibig.secondarymetabolites.org/stats)

**Figure S2.**
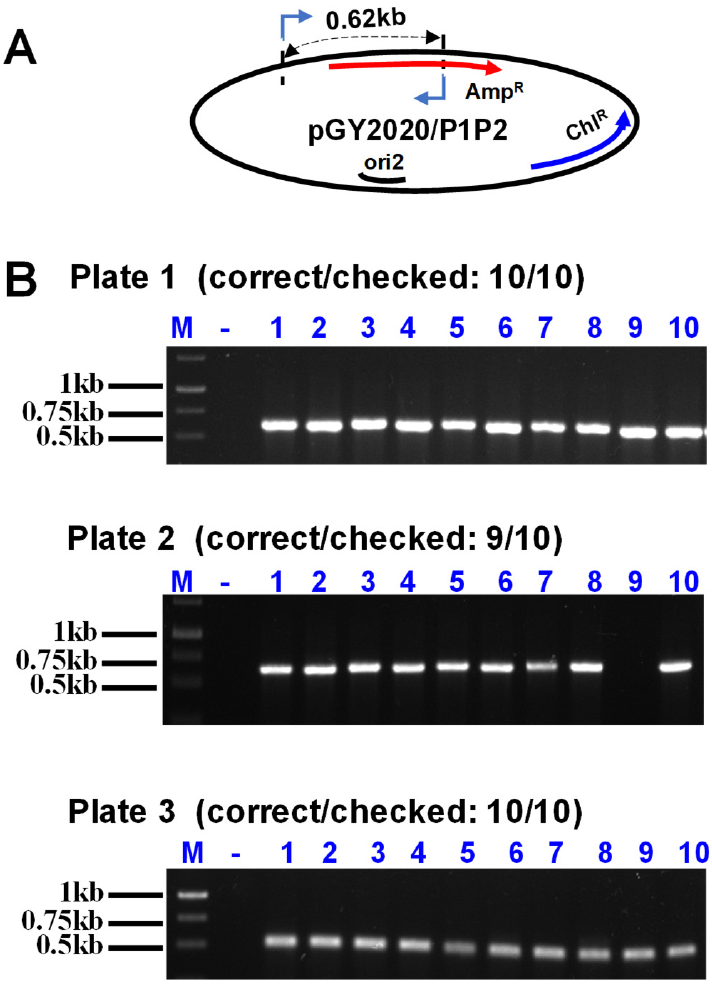
Screening of right clones by PCR amplification. A. Schematic diagram of PCR screening of right clones with primers Amp-Cas12a-scr-F/R. B. PCR amplification of ten randomly selected clones. All experiments were performed in triplicate. “-” represented blank control, genomic DNA of *E. coli* DH10B was used as PCR template.

**Figure S3.**
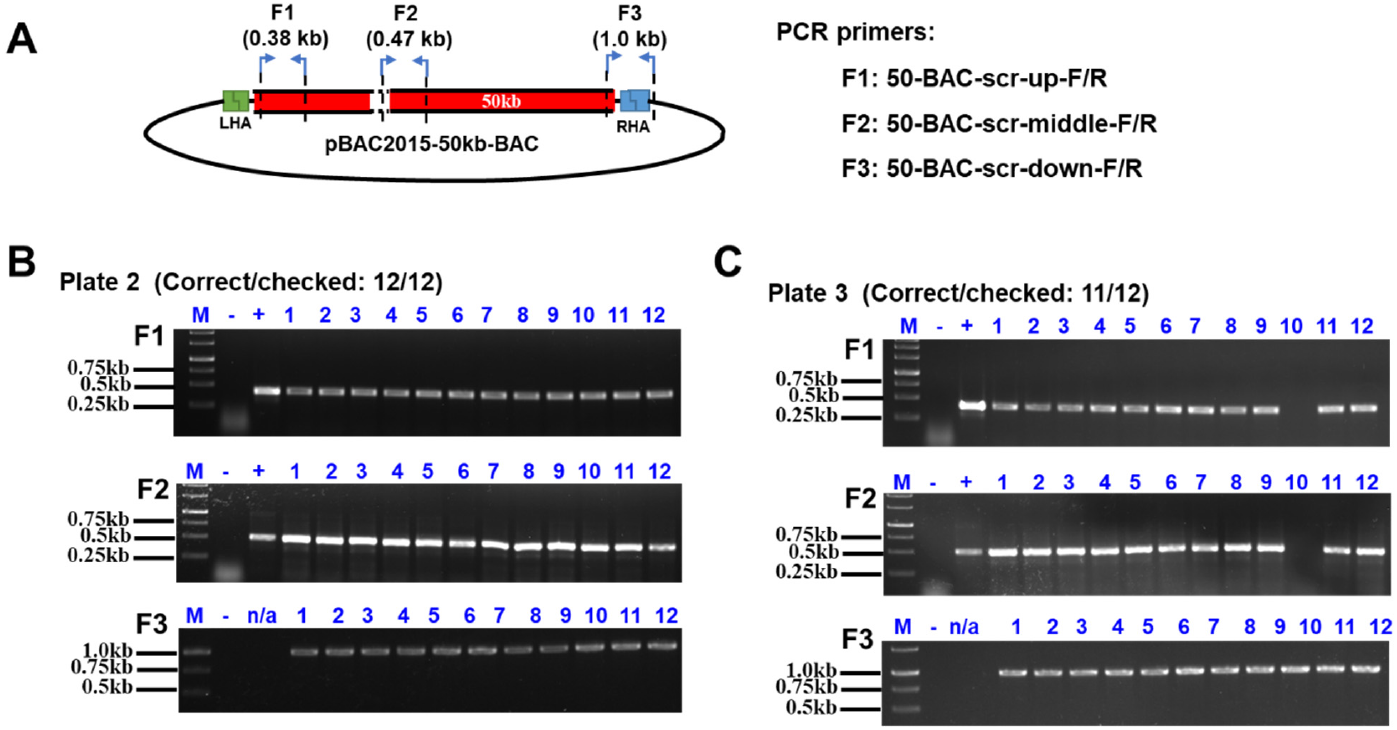
Screening of right clones that containing pBAC2015-50kb-BAC by PCR amplification. A) A. Schematic representation of crRNA design and cohesive end ligation. F1, F2 and F3 are the PCR products that were amplified using 50-BAC-scr-up-F/R, 50-BAC-scr-middle-F/R and 50-BAC-scr-down-F/R, respectively. B-C) PCR screening of 12 randomly selected clones containing pBAC2015-50kb-BAC in independent experiments/plates.

**Figure S4.**
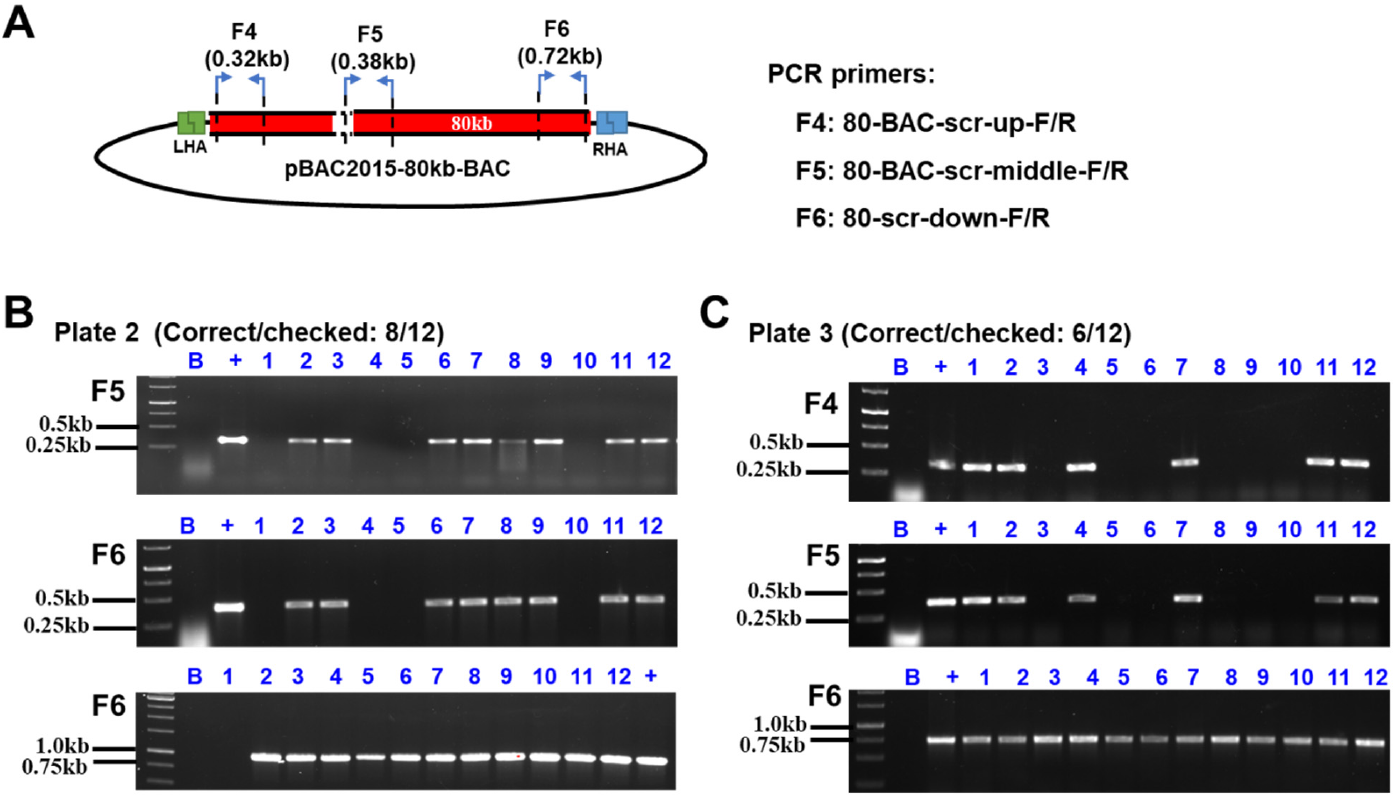
Screening of right clones that containing pBAC2015-80kb-BAC by PCR amplification. A) Schematic representation of crRNA design and cohesive end ligation. F4, F5 and F6 are the PCR products that were amplified using 80-BAC-scr-up-F/R, 80-BAC-scr-middle-F/R and 80-scr-down-F/R, respectively. Primer sequences are given in Table S7. B, C and D. PCR screening of 12 randomly selected clones containing pBAC2015-80kb-BAC in independent experiments/plates.

**Table S1.**
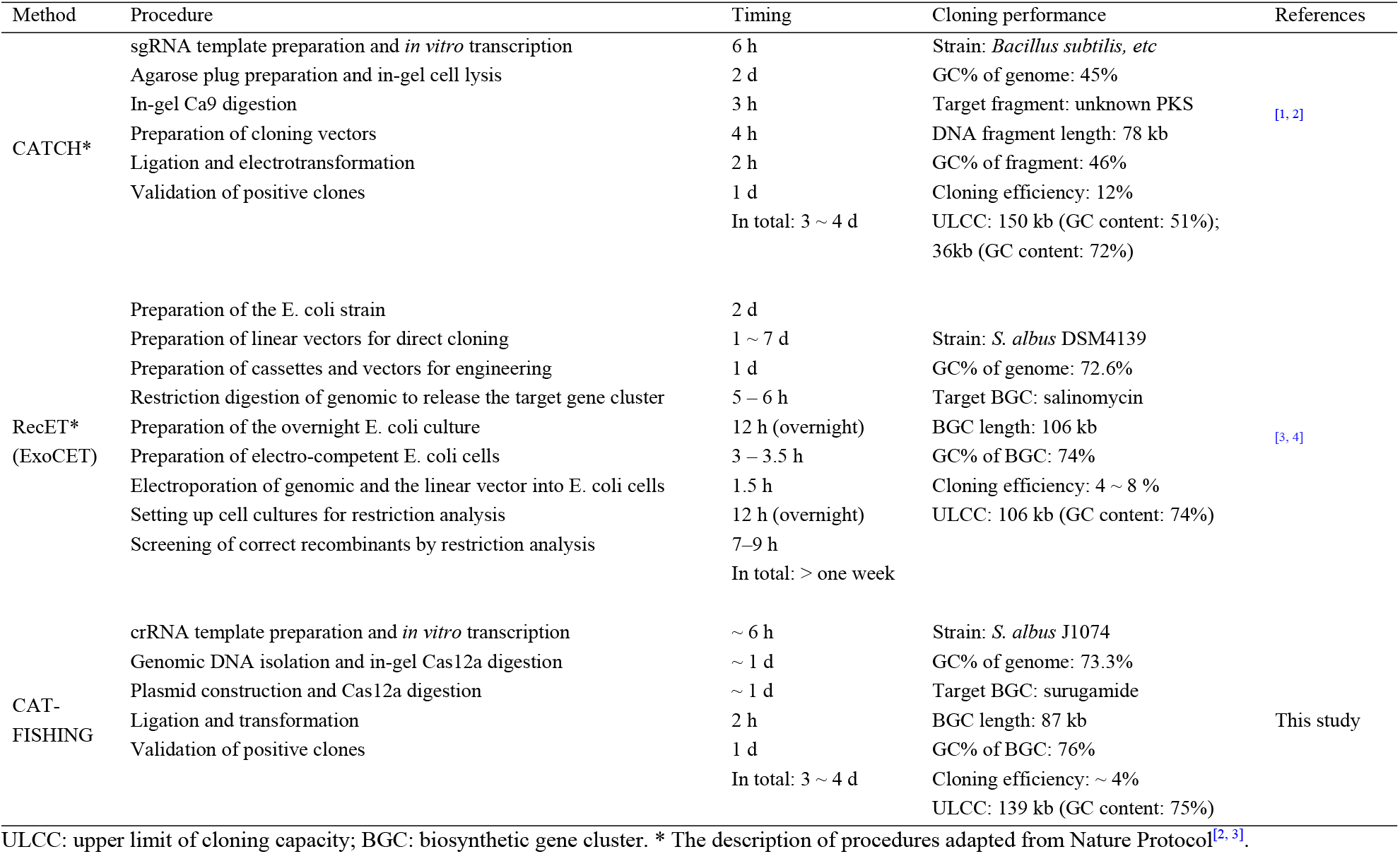
Comparison of different protocols of large DNA fragment cloning.

**Table S2.** Cost accounting of CAT-FISHING reaction system.

| Items | Consumption and cost (per 50 μL reaction) | Price & Sources |
| --- | --- | --- |
| **Cas12a** | 40 pmol; ~ 9\$ | 2000 pmol, 450\$, NEB |
| **crRNA1** | 20 pmol; ~12.5\$ | (See Table S3) |
| **crRNA2** | 20 pmol; ~12.5\$ | (See Table S3) |
| **T4 DNA ligase** | 200 U; ~ 2.2\$ | 10,000 U, 110\$, NEB |
| **Low-melt agarose** | 0.1 g; ~ 1\$ | 5 g, 30\$, Sangon Biotech |
| **β-Agarase I** | 2 U; ~2.5\$ | 100 U, 125\$, NEB |
| | | **In total: ~40\$** |

**Table S3.**
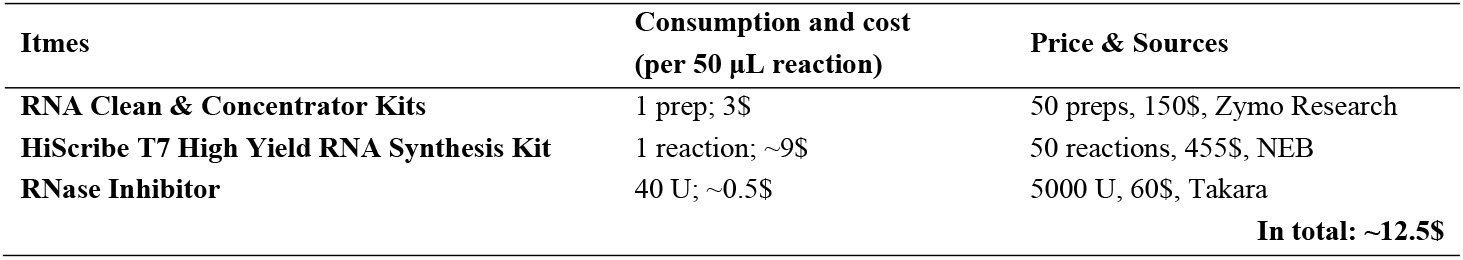
Cost accounting of crRNA preparation.

**Table S4.**
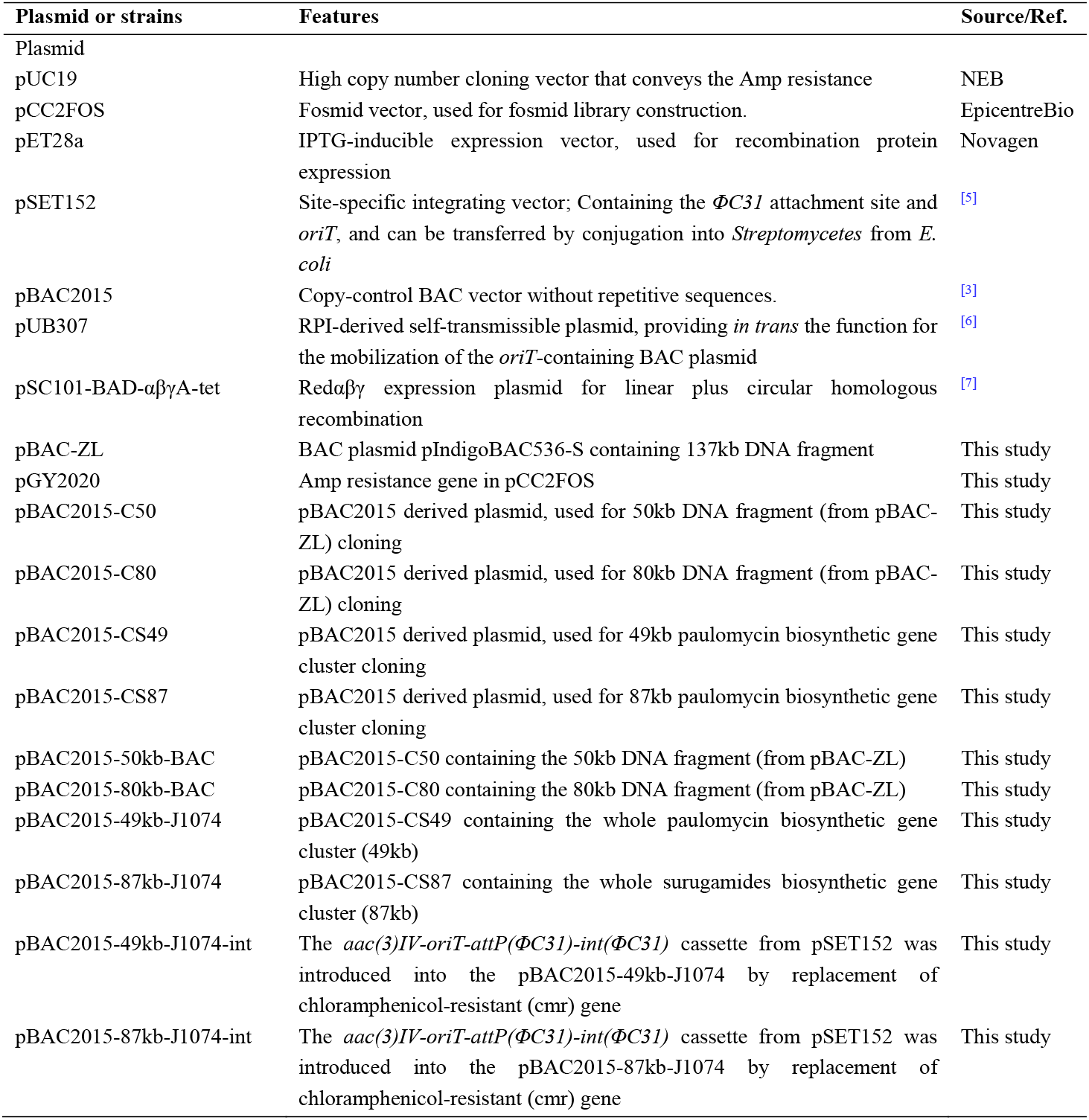
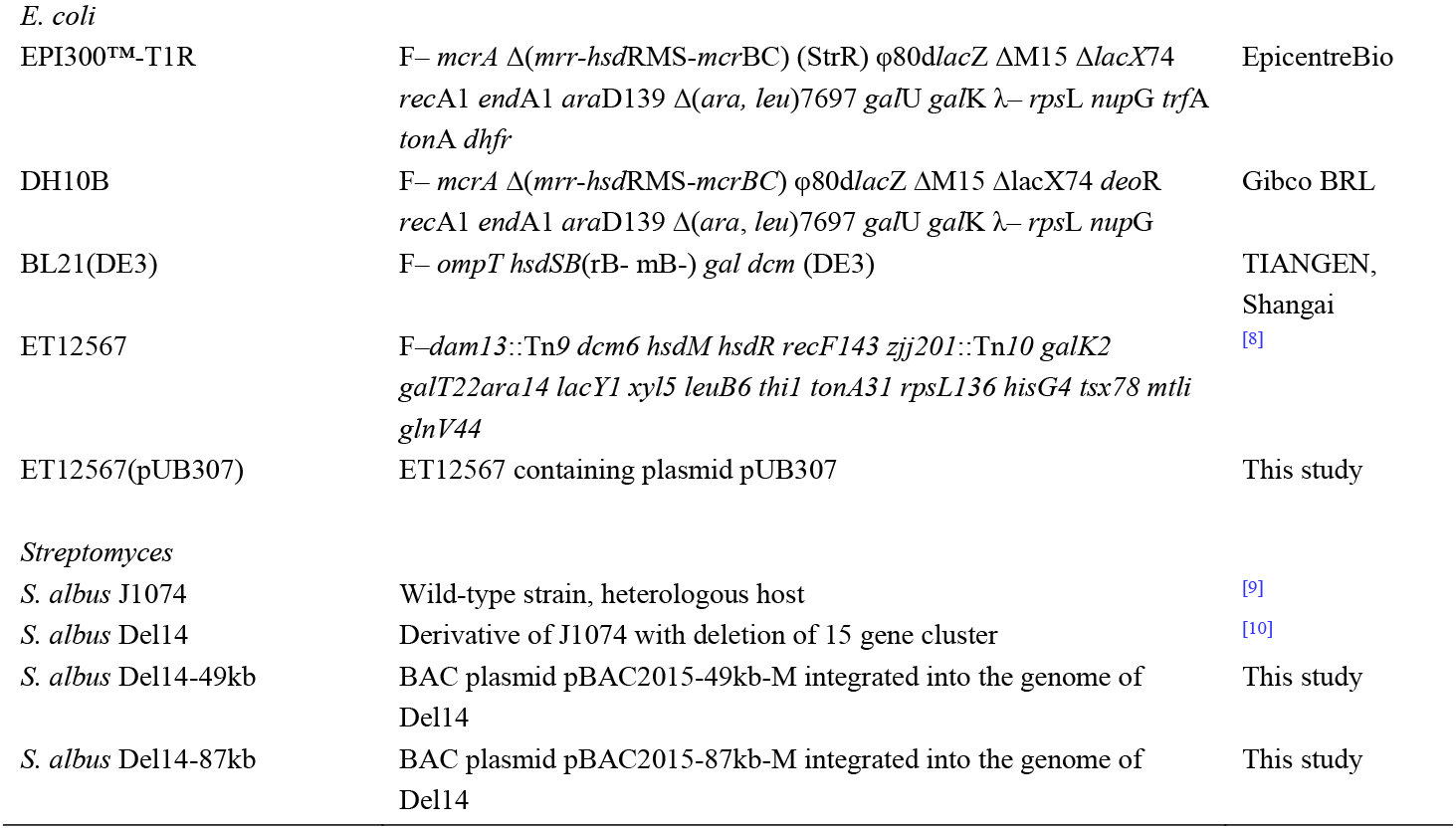
The strains and plasmids used in this study.

**Table S5.**
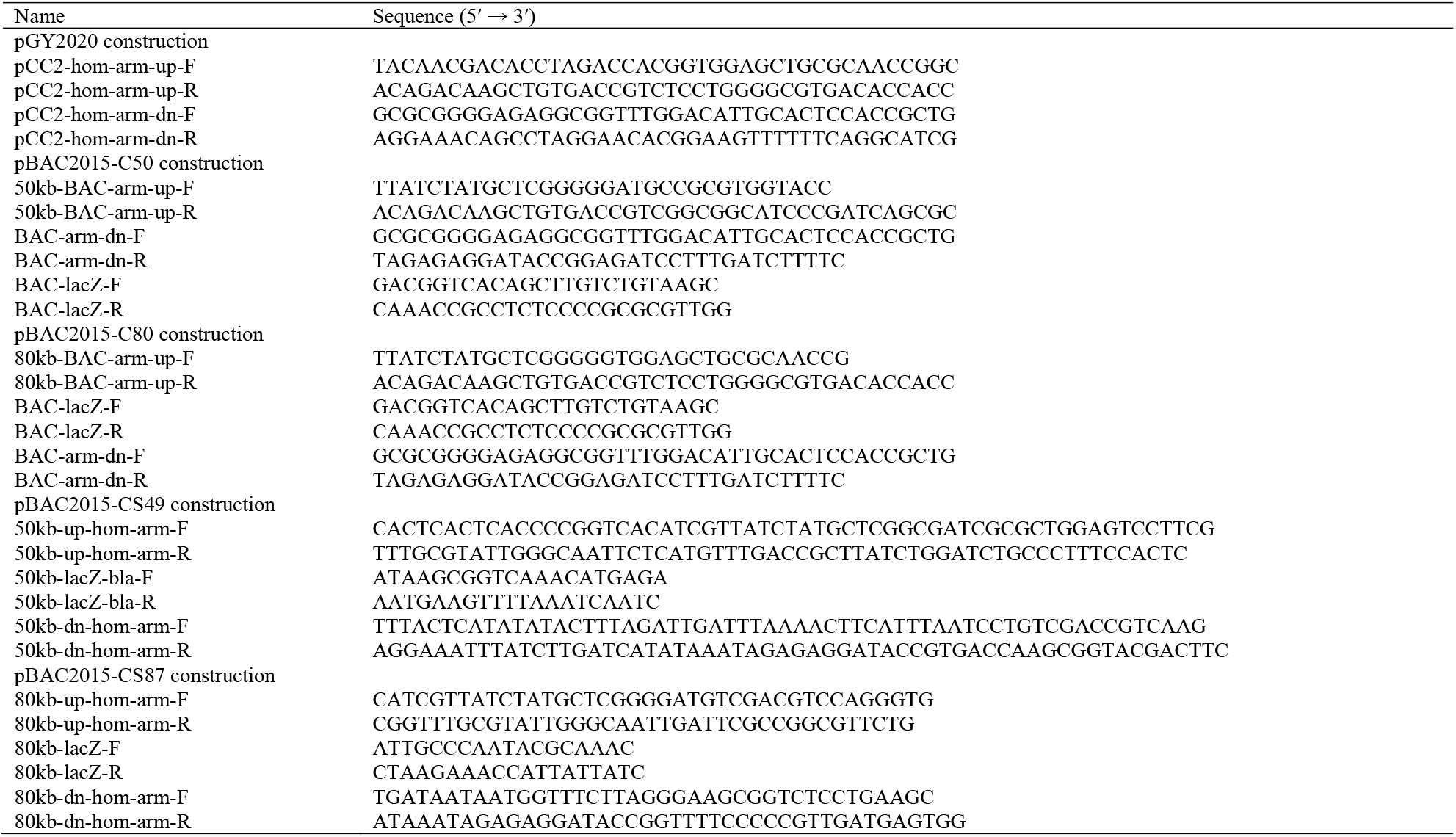
The sequence of PCR primers for plasmid construction.

**Table S6.**
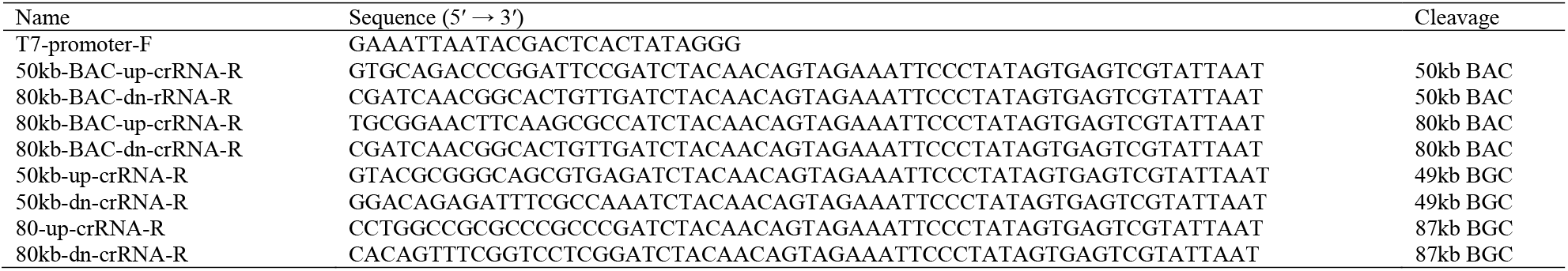
The sequence of crRNA for CRISPR/Cas12a cleavage.

**Table S7.**
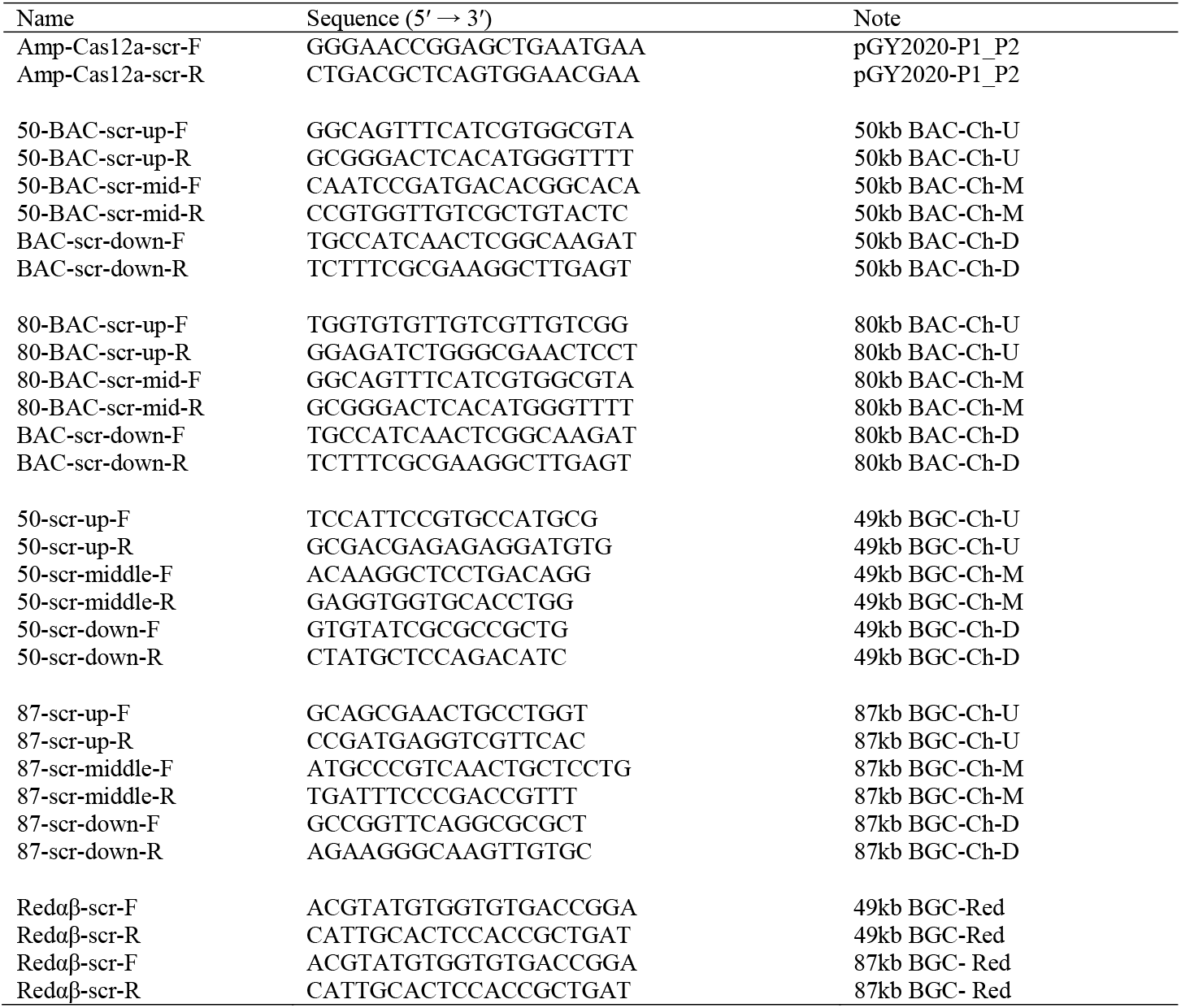
The sequence of PCR primers for screening, verification and modification.

